# Improving the Assessment of Deep Learning Models in the Context of Drug-Target Interaction Prediction

**DOI:** 10.1101/2022.04.20.488898

**Authors:** Mirko Torrisi, Antonio de la Vega de León, Guillermo Climent, Remco Loos, Alejandro Panjkovich

## Abstract

Machine Learning techniques have been widely adopted to predict drug-target interactions, a central area of research in early drug discovery. These techniques have shown promising results on various benchmarks although they tend to suffer from poor generalization. This is typically related to very sparse and nonuniform datasets available, which limits the applicability domain of machine learning techniques. Moreover, widespread approaches to split datasets (into training and test sets) treat a drug-target interaction as an independent entities, when in reality the drug and target involved may take part in other interactions, breaking apart the assumption of independence. We observe that this leads to overly optimistic test results and poor generalization of out-of-distribution samples for various state-of-the-art sequence-based machine learning models for drug-target prediction. We show that previous approaches to reduce bias in binding datasets focus on drug or target information only and, thus, lead to similar pitfalls. Finally, we propose a minimum viable solution to evaluate the generalization capability of a machine learning model based on the systematic separation of test samples with respect to drugs and targets in the training set, thus discerning the three out-of-distribution scenarios seen at test time: (1) drug or (2) target present in the training set, or (3) neither.

## 1 Introduction

Recently developed deep learning methods to predict drug-target interactions (DTI) hold great potential for drug discovery (Bagherian et al., 2021). In spite of the interest sparked both in academia and industry towards these methods, the lack of standard pipelines or benchmarking criteria discourages adoption and further developments. Particularly, as we show here, it is important to assess deep learning models beyond usual predictive measures and increase scrutiny on what the models are learning.

Data quantity and quality play a fundamental role in developing and applying deep learning models successfully. In the case of DTI, multiple databases are publicly available and focus on different characteristics of the data, such as: quality of three-dimensional structure (Binding MOAD (Smith et al., 2019)), manual curation of annotations (PDBbind (Liu et al., 2015)), or sparsity, i.e. limiting the number of drugs and targets (ExcapeDB (Sun et al., 2017)). Here, we perform an independent benchmark of DTI predictors based on BindingDB (Liu et al., 2007), a database collecting interactions from scientific articles and patents, which has already been used to create DTI benchmark datasets by others (Yingkai Gao et al., 2018; Karimi et al., 2019).

Beyond the specifics of each dataset, a crucial aspect for developing and assessing a DTI predictor is how to divide or split available data into testing and training sets, while preserving a balanced representation of the interaction space. This has been demonstrated to be particularly challenging when using chemical data for machine learning, given the amplitude of chemical space and the inherent biases present in the sparse and nonuniform DTI benchmark datasets available (Sieg et al., 2019).

Moreover, deep learning models are capable of easily fitting nuances observed in the training set, i.e. from technical noise to random labeling (Zhang et al., 2021). Although some approaches have been deployed for evaluating the redundancy between training and test sets of drug-based (Wallach & Heifets, 2018), and target-based (Urban et al., 2020) models, none of these approaches is directly applicable for DTI models. Thus, state-of-the-art DTI predictors may learn biases observed in DTI benchmark datasets instead of chemical and physical features characterizing potential interactions.

In this work, we benchmark three recently developed DTI methods and observe competitive results on two previously defined benchmark datasets. However, the models show relatively poor generalization on four Out Of Distribution (OOD) scenarios. Intuitively, a DTI deep learning model will generalize to OOD scenarios by learning fundamental chemical and physical features that govern the interaction between a drug and a target at the molecular level. To further understand the generalization challenge, we investigated if considering interactions as independent entities, as done previously when splitting the training and test samples in these defined benchmark datasets, may constitute a potential source of information leakage which, in turn, may hinder the learning and evaluation of DTI predictors. With a baseline analysis, we show that the association of DTI in these benchmark datasets by drug or target alone contains information and predictive power and, thus, constitutes a case of information leakage.

To improve the assessment of DTI predictors, we propose a minimum viable solution for measuring their generalization capability, i.e. discerning the three OOD scenarios seen at test time: (1) drug or (2) target present in the training set, or neither (3). Our solution strengthens the assessment of DTI predictors evaluating empirically their ability to generalize to OOD scenarios, without altering the training set, while facilitating the identification of potential sources of bias in benchmark datasets.

To summarize, we show that considering DTI as independent entities introduces potential sampling biases which may hinder the prediction and generalization capability of a DTI model. Correspondingly, we propose a simple approach to gauge the generalization capability of a DTI model emphasizing the evaluation on OOD scenarios.

## 2 Methods

### 2.1 Datasets

All the datasets in this work are derived from BindingDB (Liu et al., 2007): a publicly accessible and regularly updated collection of binding affinity values between proteins considered to be drug-targets, and drug-like molecules. In particular, we adopt two benchmark datasets derived from BindingDB, one released by Yingkai Gao et al. (2018) and the other as defined by Karimi et al. (2019), which have been used for benchmarking recent DTI predictors (Chen et al., 2020; Born et al., 2022). Both benchmark datasets are outlined in Table 1.

**Table 1:** Number of unique drugs, targets, and interactions observed in the benchmark datasets, i.e. *BindingDB^S^* and *BindingDB^L^*, as well as sparsity and ratio of Positive/Negative samples. Values within parenthesis are the overlap with the training set, omitted when equal to 0%, bolded when greater than 50%, and underlined in cases of data leakage.

| Dataset | Drugs | Targets | Interactions | Sparsity | P/N |
| --- | --- | --- | --- | --- | --- |
| *train<sup>S</sup>* | 41,708 | 741 | 48,512 | 0.16% | 1.26 |
| *val<sup>S</sup>* | 4,882 (30%) | 456 (**90%**) | 5,376 | 0.24% | 1.02 |
| *test<sup>S</sup>* | 4,798 (30%) | 455 (**90%**) | 5,248 | 0.24% | 0.97 |
| Total (unique entities) | 48,084 | 798 | 59,136 | 0.15% | 1.21 |
| *train<sup>L</sup>* | 180,736 | 2,632 | 232,044 | 0.05% | 1.30 |
| *val<sup>L</sup>* | 25,215 (39%) | 1,778 (**98%**) | 26,272 (4% ) | 0.06% | 1.33 |
| *test<sup>L</sup>* | 97,358 (37%) | 2,370 (**95%**) | 111,906 (<u>4%</u>) | 0.05% | 1.31 |
| *test<sup>Ch</sup>* | 12,795 (18%) | 125 (1%) | 14,107 | 0.88% | 1.02 |
| *test<sup>ER</sup>* | 2,115 (13%) | 6 | 3,228 | 25% | 1.5 |
| *test<sup>GPCR</sup>* | 48,712 (13%) | 313 | 57,957 | 0.38% | 2.74 |
| *test<sup>RTK</sup>* | 24,608 (25%) | 127 (1%) | 33,189 | 1.06% | 3.31 |
| Total (unique entities) | 321,950 | 3,350 | 472,925 | 0.05% | 1.48 |

The first version, which we name *BindingDB^S^*, contains 59,136 interactions (involving 48,084 drugs and 798 targets), distributed in training (*train^S^*), validation (*val^S^*), test (*test^S^*) set. It was assembled with the intent of simulating practical scenarios, i.e. “given a pair of drug and target at testing time, the drug, the target, or both of them may have not been observed at training time” (Yingkai Gao et al., 2018). We remove from the original sets any drug and target exceeding the length constraints of DeepAffinity (Karimi et al., 2019) (see Section 2.2).

The second version, *“BindingDB^L^*”, contains 472,925 interactions (involving 321,950 drugs and 3,350 targets), distributed in training (*train^L^*), validation (*val^L^*), test (*test^L^*), and 4 OOD sets, i.e. ion Channels (*test^Ch^*), nuclear Estrogen Receptors (*test^ER^*), G-Protein-Coupled Receptors (*test^GPCR^*), and Receptor Tyrosine Kinases (*test^RTK^*). This collection of datasets was released for assessing various deep learning techniques and their generalization capability, constraining four protein families only in the respective OOD sets (Karimi et al., 2019), i.e. leaving ion Channels, nuclear Estrogen Receptors (*test^ER^*), G-Protein-Coupled Receptors, and Receptor Tyrosine Kinases out of the training set. Although none of the method in this work needs a validation set, we follow the approach used by the authors of DeepAffinity (Karimi et al., 2019), and randomly select 10% of the samples in the training set to split the validation set.

### 2.2 DTI Predictors

In this work, we assess the predictive performance of three recent deep learning methods, together with a machine learning baseline: DeepAffinity (Karimi et al., 2019), DeepConv-DTI (Lee et al., 2019), TransformerCPI (Chen et al., 2020), and Random Forest (Ho, 1995). All deep learning methods are published along with the code for training and assessing them. We do not perform any hyperparameter optimization and aim to reproduce the methods as described in the respective manuscripts.

DeepAffinity (Karimi et al., 2019) is the only regressor in this work, which makes comparison to the other methods challenging. However, for completeness, we include comparison with this method. To do this, we impose a classification threshold of 0.5 for *BindingDB^S^* (which was released already binarized), and of 6 for *BindingDB^L^* (as done previously on *BindingDB^S^* by Yingkai Gao et al. (2018)). DeepAffinity requires the PDB (Berman et al., 2000), Pfam (Mistry et al., 2021) and UniRef (Suzek et al., 2015) for building the vocabulary of 72 four-mers describing the targets. Drugs are described using an alphabet of 68 letters derived from the corresponding SMILES. Moreover, UniRef50 (Suzek et al., 2015) and Stitch (Szklarczyk et al., 2016) are used for pre-training the embeddings of drugs and targets, respectively. DeepAffinity accepts targets of up to 1,500 amino acids, and SMILES of up to 100 symbols.

DeepConv-DTI (Lee et al., 2019) is the only method not relaying on external databases, nor k-mers, nor pre-training. Although it relays on an embedding for representing protein sequences, such embedding is trained from scratch on the training set. Drugs are represented by Morgan fingerprints of radius 2 (Rogers & Hahn, 2010). It accepts targets of up to 2,500 amino acids.

TransformerCPI (Chen et al., 2020) is inspired from a Transformer architecture (Vaswani et al., 2017), although the drugs are passed in input only to the decoder. Protein sequences are split into overlapping three-mers of amino acids, and then passed to a word2vec embedding (Mikolov et al., 2013) (pre-trained on human proteins from UniProt (The UniProt Consortium, 2017)). Each atom of the drugs is represented via 34 chemical and physical properties calculated with RDKit (RDKit), which are then passed to a graph.

Random Forest (Ho, 1995) is implemented using the code released along with DeepAffinity (Karimi et al., 2019). In particular, protein sequences are represented by a vector of size 2,500 (padded with zeros for shorter sequences), where each amino acid is encoded via a label encoder, i.e. mapped to a number from one to twenty-five. Drugs are represented using Morgan fingerprints of radius 2 (Rogers & Hahn, 2010) calculated with RDKit (RDKit).

### 2.3 Metrics

We use Matthews Correlation Coefficient (MCC) to measure predictive performance. MCC summarizes the confusion matrix in a single value, and is considered to be more informative than other metrics on imbalanced datasets (Brown, 2018). Specifically, the MCC can be calculated from the confusion matrix as follow:

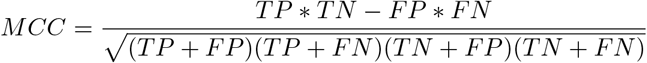

with TP (True Positives), TN (True Negatives), FP (False Positives), and FN (False Negatives). The MCC can assume values from −1 to 1, where the extremes indicate perfect agreement or disagreement between ground truth and model predictions, and 0 indicates no relationship.

For completeness, we include Accuracy and F1 metrics for all results in this work in the AppendixA.

## 3 Results

### 3.1 DTI Benchmark

We benchmark DeepAffinity, DeepConv-DTI, TransformerCPI, and a Random Forest on *BindingDB^S^* and *BindingDB^L^*. To do so, we train all methods twice to obtain a model for each benchmark dataset.

We observe good performance across the board on *test^S^* and *test^L^*, as shown in Figure 1a. DeepConv-DTI achieves a MCC over 0.7, slightly outperforming other methods. Notably, the Random Forest is never the worst method in this setting, and appears to be on par with the deep learning methods.

**Figure 1:**
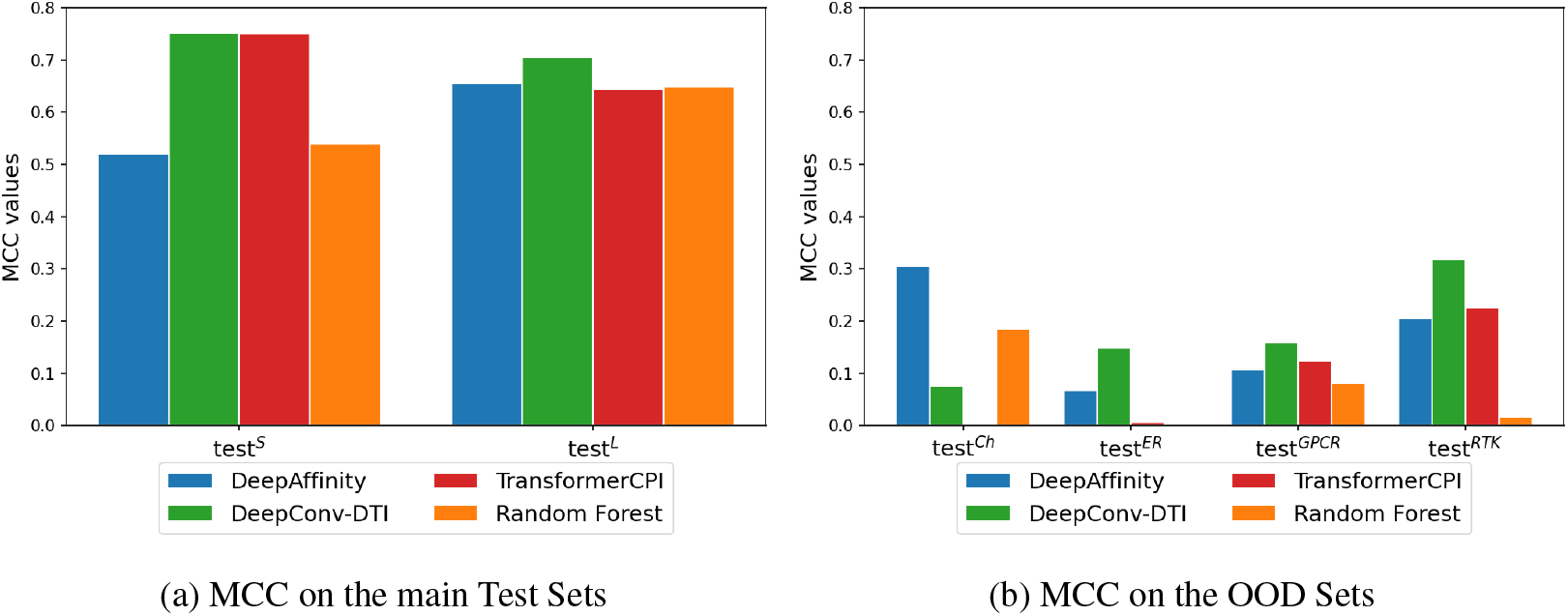
MCC observed on (a) *test^S^* and *test^L^*, and on (b) the four OOD sets. All methods provide good predictive performance on the test sets, and lower performance on the OOD sets.

Lower performance is observed regarding generalization, as measured on the four OOD sets, i.e. *test^Ch^, test^ER^, test^GPCR^*, and *test^RTK^* (see Figure 1b). All methods achieve lower MCC values than on *test^S^* and *test^L^*, and exceed a MCC value of 0.3 only in two cases, i.e. DeepAffinity on *test^Ch^*, and DeepConv-DTI on *test^RTK^*. DeepConv-DTI achieves the highest MCC values in most cases, although no method outperforms the competitors on all four OOD sets, and the Random Forest is generally the worst by MCC value.

The results of our benchmark outline that no method outperforms all the other methods assessed, in all benchmarking scenarios. Importantly, all methods achieve good MCC on *test^S^* and *test^L^*, but much lower performance on OOD scenarios, pointing towards poor generalization. Regarding DeepAffinity, we observe particularly good predictive performance on *test^L^* and *test^Ch^*, although it is challenging to compare the only regressor with the other methods.

### 3.2 Drugs and Targets distribution

To identify possible sources of overoptimistic performance on *test^S^* and *test^L^*, which may justify the poor generalization on *test^Ch^, test^ER^, test^GPCR^*, and *test^RTK^*, we look deeper in the composition of the benchmark datasets. In Section 3.2.1, we explore the overlap between training and testing sets, first at the level of interactions and then at the level of their constitutive drugs and targets. In Section 3.2.2, we performed a baseline analysis to measure the potential information leakage because of the overlap of drugs and targets across different interactions between training and testing sets. Our results show that treating DTI as independent entities causes information leakage in the context of train-test splits.

#### 3.2.1 Overlaps across sets

We find overlapping interactions in *BindingDB^L^*, a simple case of data leakage. Specifically, we observe that 4% of the interactions in both *test^L^* and *val^L^* are present in *train^L^* (see Table 1). We also observe DTI with multiple labels, i.e. the same drug-target pair associated to multiple binding observations. This case of data leakage may be due to the lack of stereochemistry in the SMILES of *BindingDB^L^*, leading different molecules to be treated as if they were the same.

Beyond direct overlap of interactions between datasets, we also look at interactions not as independent entities, but in terms of their constitutive drugs and targets (see Table 1). In this regard, we find overlap of drugs and targets across all datasets with respect to their training set, including *test^Ch^, test^ER^, test^GPCR^*, and *test^RTK^*.

We find that 30% of drugs and 90% of targets in *test^S^* are also in *train^S^*, as shown in Table 1. We find an even greater overlap between *test^L^* and *train^L^*, i.e. 37% of drugs and 95% of targets. Similar overlaps exist also between the validation sets and the respective training sets for both *BindingDB^S^* and *BindingDB^L^*, as shown in Figure 2. In the case of the four OOD sets, we find that 13-25% of the drugs in these sets, as well as one target in *test^Ch^* and a second one in *test^RTK^*, are present in *train^L^*.

**Figure 2:**
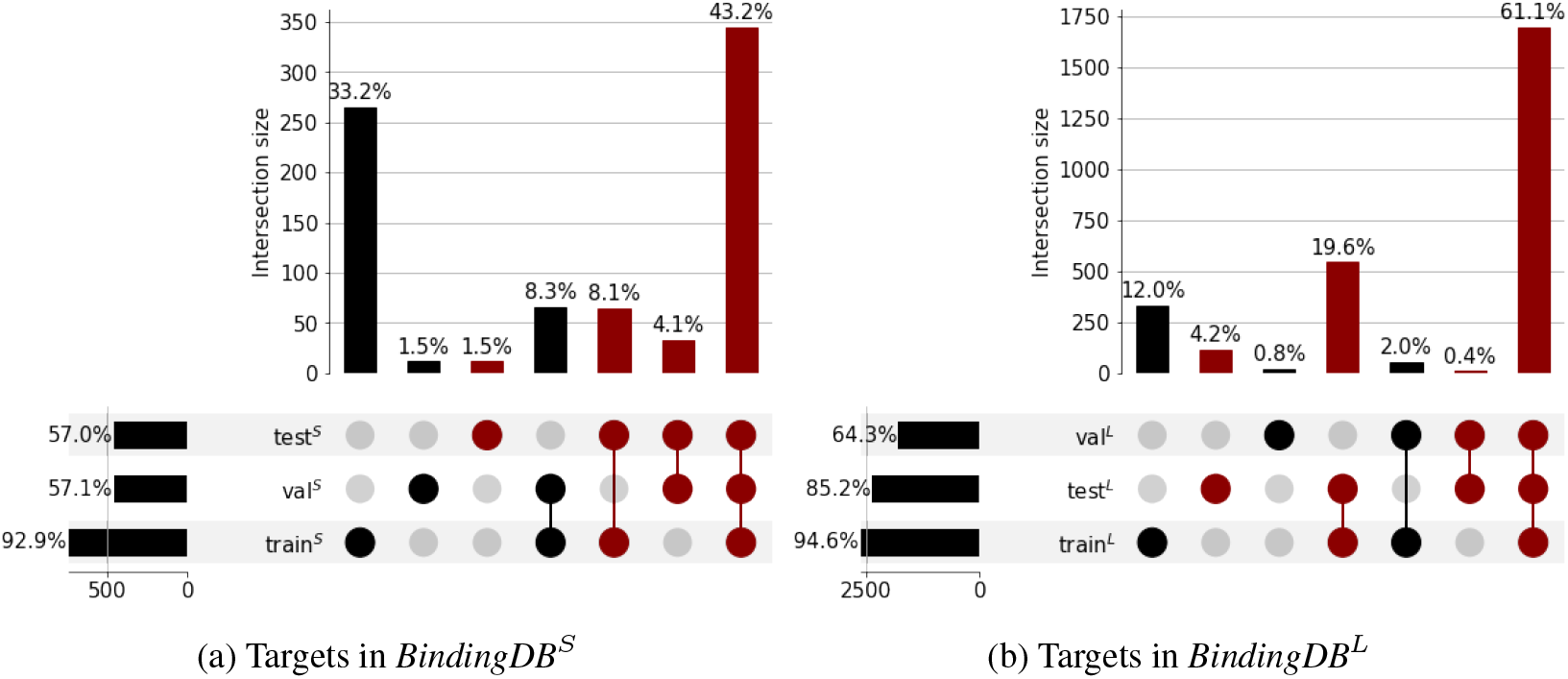
Target overlaps in (a) *BindingDB^S^* and in (b) *BindingDB^L^* with respect to the entire benchmark datasets. In both cases, a large portion of targets are present in training, validation, and test set at the same time.

We further investigate the large overlaps of targets across training, validation, and test set seen in *BindingDB^S^* and *BindingDB^L^*, looking at the most represented targets. In particular, we pick the top 10% targets by number of interactions in each dataset, i.e. considering the number of interactions in each dataset individually. We observe that nearly all top targets in *test^S^* and *test^L^* are among the top targets in the respective training and validation set, as shown in Figure 3. Thus, the overlap of targets across training, validation, and test set seen in *BindingDB^S^* and *BindingDB^L^* is strongly present even when looking at the top targets.

**Figure 3:**
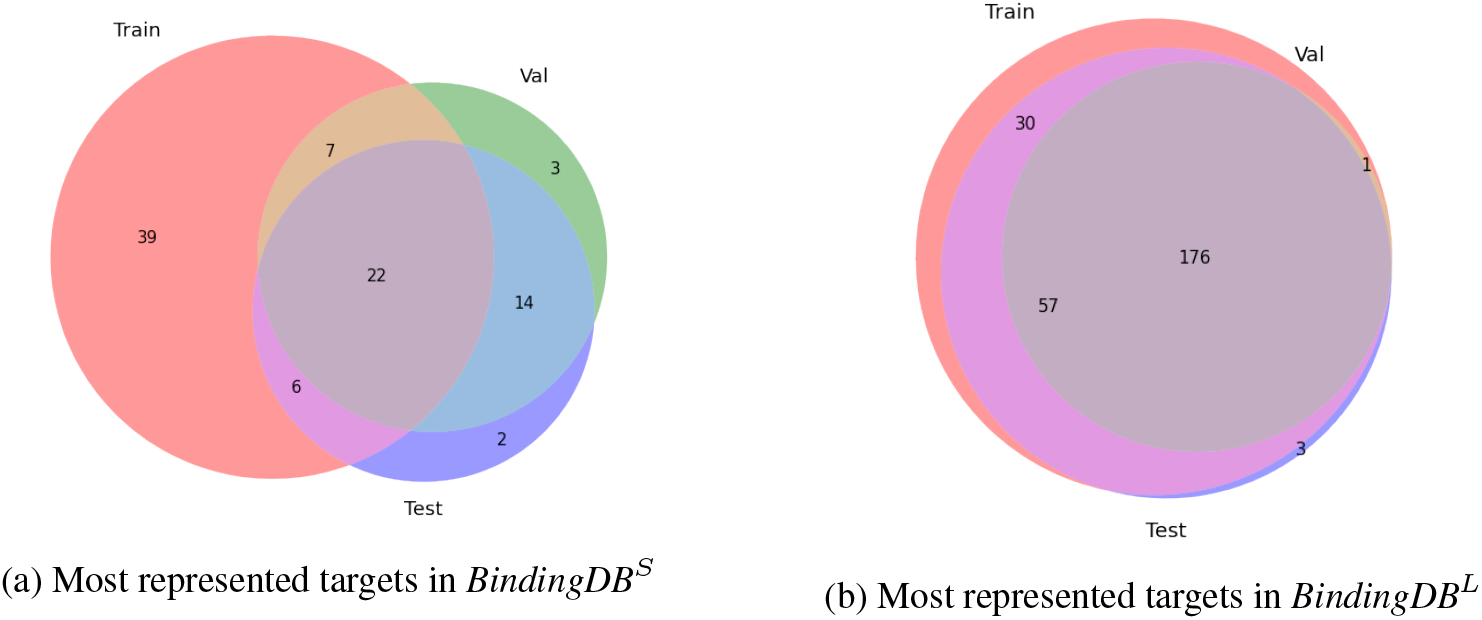
The overlap of targets across training, validation, and test set is strongly present even when looking at the subset of the most represented targets in (a) *BindingDB^S^* and in (b) *BindingDB^L^*. The overlap between training and test set is shown in magenta, and in grey when overlapping also the validation set.

#### 3.2.2 Identity matching

The overlaps of drugs and targets we observe across the benchmark datasets may represent a problematic case of information leakage, which may hinder the learning of DTI model.

We investigate this hypothesis performing a baseline analysis by extracting the average activity value for each drug and target in the training set. Then, for each interaction in the test set, if it contains a drug or target that is present in the training set, a drug-based prediction (p_drug) and/or a target-based prediction (p_target) is provided, corresponding to their average activity in the training set.

This baseline analysis matches identities for drugs and targets across training and test sets; it performs no machine learning, does not access any protein sequence or compound chemical information, nor does it use any similarity or clustering calculation. Thus, this baseline cannot analyze interactions involving a drug and a target when both are not present in the training set (see Table 6 for the number of predictions for each dataset). Given the lack of chemical and protein sequence information in this approach, one would expect a performance close to random (MCC close to 0.0). Nonetheless, this approach was able to produce surprisingly high MCC values on both *BindingDB^S^* and *BindingDB^L^*, as shown in Figure 4b (see full results in Table 6).

**Figure 4:**
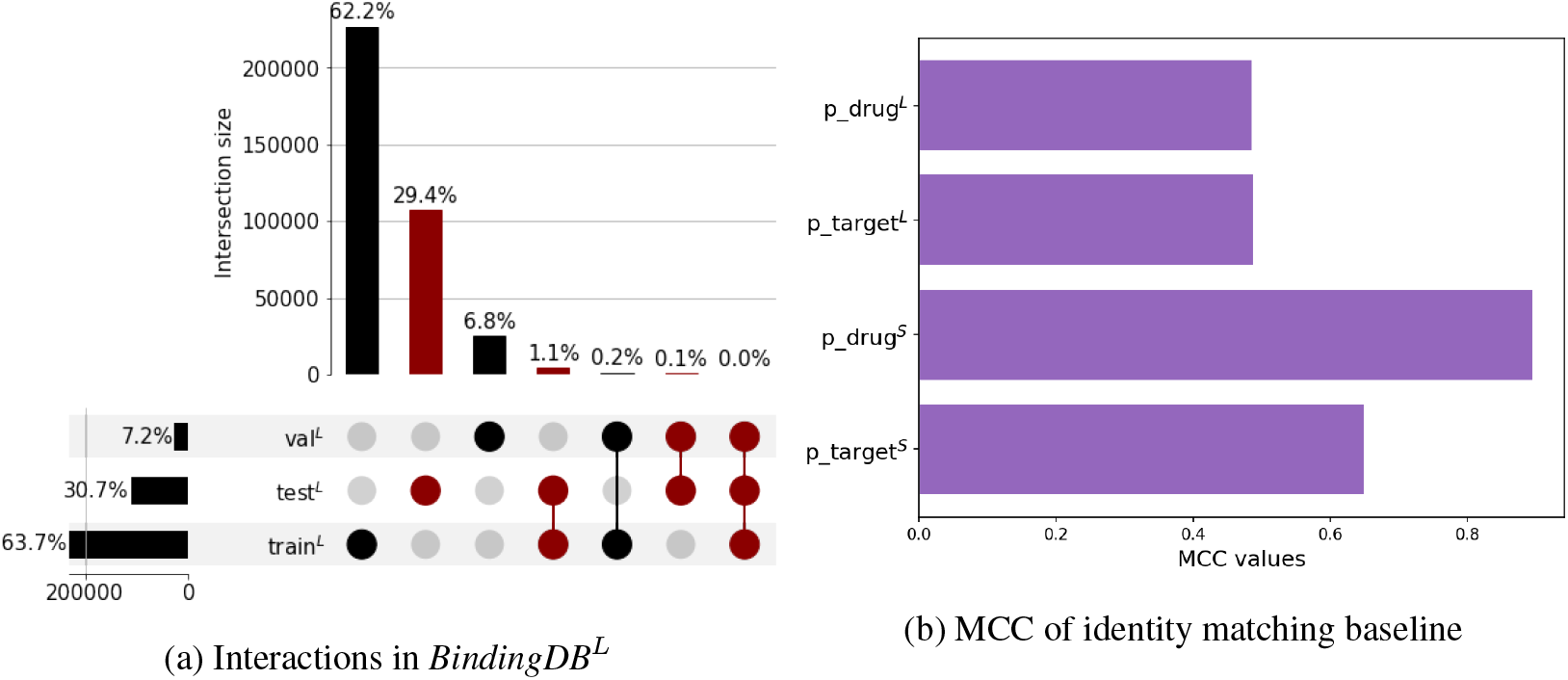
(a) Overlaps of interactions in *BindingDB^L^* with respect to the entire benchmark dataset; (b) baseline analysis matching identities for drugs and targets across training and test sets: for each interaction in the test set, a drug-based prediction (p_drug) and/or a target-based prediction (p_target) is provided, corresponding to their average activity in the training set.

Therefore, this analysis provides evidence of the information leakage present in the dataset due to the non-independent nature of interactions with respect to their constitutive drugs and targets.

### 3.3 Evaluation of Generalization Capability

Motivated by the information leakage we find in *BindingDB^S^* and *BindingDB^L^*, we investigate whether filtering the test samples according to the overlap with the training set provides a more informative evaluation of the generalization capability of a method. Therefore, we propose to disaggregate OOD scenarios to improve the benchmarking of DTI predictors.

In particular, we derive 3 subsets for each test set collecting any interaction where:

1. the drug is not in the training set (*D*);
2. the target is not in the training set (*T*);
3. neither the target nor the drug is in the training set (*I*).

In the Sections below, we use this approach to shed additional light on the information leakage that partially causes the overoptimistic predictive performance observed on *test^S^* and *test^L^*.

#### 3.3.1 BindingDB^S^

The filtering approach we propose results in three subsets of *test^S^*, i.e. *D^S^, T^S^*, and *I^S^* (see Table 2). For all methods, we observe good predictive performance on *test^S^* and *D^S^*, as shown in Figure 5a. Lower performance is shown on *T^S^* and *I^S^*, meaning that filtering by target has a similar effect to filtering by both drug and target in this dataset. The stronger relevance of filtering by target matches the larger overlap seen across targets for *D^S^* (as show in Table 2). In part, this can be expected due to the composition of the data (one order of magnitude fewer targets than drugs), but it also points towards the information leakage present in *test^S^*, due to the non-independent nature of drugs and targets within interactions.

**Table 2:** Number of unique drugs, targets, and interactions observed in the modified benchmark datasets, as well as sparsity and ratio of Positive/Negative samples. Values within parenthesis are the overlap with the training set, omitted when equal to 0%, and bolded when greater than 50%.

| Dataset | Drugs | Targets | Interactions | Sparsity | P/N |
| --- | --- | --- | --- | --- | --- |
| $D^S$ | 3,354 | 356 (**90%**) | 3,623 | 0.3% | 1.33 |
| $T^S$ | 2,210 (15%) | 45 | 2,357 | 2.36% | 1.03 |
| $I^S$ | 1,890 | 35 | 2,001 | 3.03% | 1.36 |
| Total (unique entities) | 3,674 | 366 | 3,979 | 2.41% | 1.13 |
| $train^R$ | 195,479 | 2,440 | 265,510 | 0.06% | 1.3 |
| $val^R$ | 28,170 (42%) | 1,678 (**97%**) | 29,501 | 0.06% | 1.29 |
| $test^R$ | 57,250 (22%) | 1,996 (**85%**) | 69,433 | 0.06% | 1.21 |
| $D^R$ | 35,268 | 1,945 (**87%**) | 54,662 | 0.34% | 1.14 |
| $T^R$ | 43,594 (48%) | 305 | 36,694 | 0.07% | 1.26 |
| $I^R$ | 21,612 | 254 | 21,923 | 0.4% | 1.22 |
| $I^{Ch}$ | 9,917 | 116 | 11,016 | 0.96% | 1.13 |
| $I^{ER}$ | 1,779 | 6 | 2,773 | 25.98% | 1.29 |
| $I^{GPCR}$ | 40,606 | 272 | 46,746 | 0.42% | 1.8 |
| $I^{RTK}$ | 12,254 | 93 | 21,255 | 1.33% | 3.99 |
| Total (unique entities) | 321,891 | 3,268 | 446,234 | 0.04% | 1.46 |

**Table 3:**
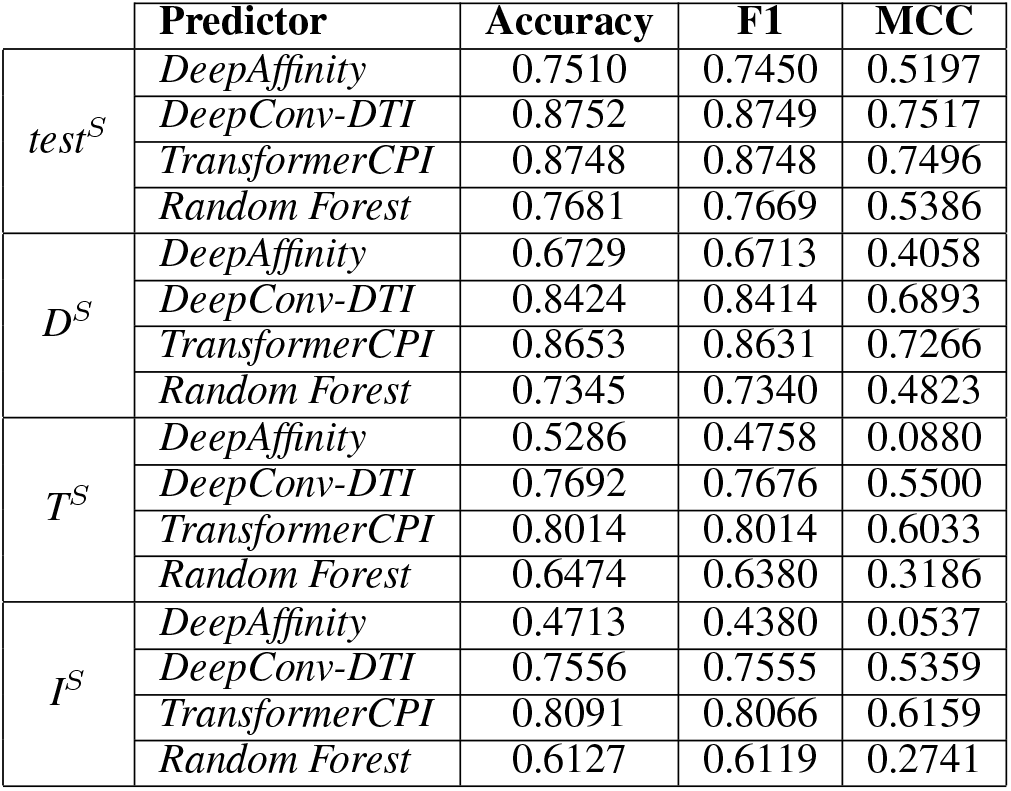
Results of DTI benchmark on the test set and subsets of *BindingDB^S^*.

|  | Predictor | Accuracy | F1 | MCC |
| --- | --- | --- | --- | --- |
| $test^S$ | *DeepAffinity* | 0.7510 | 0.7450 | 0.5197 |
|  | *DeepConv-DTI* | 0.8752 | 0.8749 | 0.7517 |
|  | *TransformerCPI* | 0.8748 | 0.8748 | 0.7496 |
|  | *Random Forest* | 0.7681 | 0.7669 | 0.5386 |
| $D^S$ | *DeepAffinity* | 0.6729 | 0.6713 | 0.4058 |
|  | *DeepConv-DTI* | 0.8424 | 0.8414 | 0.6893 |
|  | *TransformerCPI* | 0.8653 | 0.8631 | 0.7266 |
|  | *Random Forest* | 0.7345 | 0.7340 | 0.4823 |
| $T^S$ | *DeepAffinity* | 0.5286 | 0.4758 | 0.0880 |
|  | *DeepConv-DTI* | 0.7692 | 0.7676 | 0.5500 |
|  | *TransformerCPI* | 0.8014 | 0.8014 | 0.6033 |
|  | *Random Forest* | 0.6474 | 0.6380 | 0.3186 |
| $I^S$ | *DeepAffinity* | 0.4713 | 0.4380 | 0.0537 |
|  | *DeepConv-DTI* | 0.7556 | 0.7555 | 0.5359 |
|  | *TransformerCPI* | 0.8091 | 0.8066 | 0.6159 |
|  | *Random Forest* | 0.6127 | 0.6119 | 0.2741 |

**Table 4:** Results of DTI benchmark on the test set and OOD sets of *BindingDB^L^*.

|  | Predictor | Accuracy | F1 | MCC |
| --- | --- | --- | --- | --- |
| $test^L$ | *DeepAffinity* | 0.8317 | 0.8279 | 0.6560 |
|  | *DeepConv-DTI* | 0.8559 | 0.8525 | 0.7055 |
|  | *TransformerCPI* | 0.8259 | 0.8215 | 0.6439 |
|  | *Random Forest* | 0.8279 | 0.8219 | 0.6480 |
| $test^{Ch}$ | *DeepAffinity* | 0.6468 | 0.6435 | 0.3052 |
|  | *DeepConv-DTI* | 0.5372 | 0.5367 | 0.0767 |
|  | *TransformerCPI* | 0.4961 | 0.4953 | -0.0091 |
|  | *Random Forest* | 0.5420 | 0.4509 | 0.1858 |
| $test^{ER}$ | *DeepAffinity* | 0.4899 | 0.4879 | 0.0671 |
|  | *DeepConv-DTI* | 0.5101 | 0.5029 | 0.1482 |
|  | *TransformerCPI* | 0.4988 | 0.4942 | 0.0063 |
|  | *Random Forest* | 0.3856 | 0.2783 | 0.0000 |
| $test^{GPCR}$ | *DeepAffinity* | 0.6245 | 0.5501 | 0.1060 |
|  | *DeepConv-DTI* | 0.6022 | 0.5615 | 0.1596 |
|  | *TransformerCPI* | 0.6261 | 0.5577 | 0.1240 |
|  | *Random Forest* | 0.3884 | 0.3860 | 0.0808 |
| $test^{RTK}$ | *DeepAffinity* | 0.5516 | 0.5304 | 0.2060 |
|  | *DeepConv-DTI* | 0.7150 | 0.6475 | 0.3173 |
|  | *TransformerCPI* | 0.6853 | 0.6051 | 0.2263 |
|  | *Random Forest* | 0.2397 | 0.2070 | 0.0159 |

**Table 5:** Results of DTI benchmark on the test set, subsets, and OOD sets of *BindingDB^R^*.

|  | Accuracy | F1 | MCC |
| --- | --- | --- | --- |
| $test^R$ | 0.6894 | 0.6890 | 0.4023 |
| $D^R$ | 0.7040 | 0.7040 | 0.4227 |
| $T^R$ | 0.5631 | 0.5389 | 0.1883 |
| $I^R$ | 0.5729 | 0.5441 | 0.1920 |
| $I^{Ch}$ | 0.4971 | 0.3952 | 0.1047 |
| $I^{ER}$ | 0.3567 | 0.2629 | 0.0000 |
| $I^{GPCR}$ | 0.3636 | 0.3606 | 0.0520 |
| $I^{RTK}$ | 0.2031 | 0.1719 | -0.0173 |

**Table 6:**
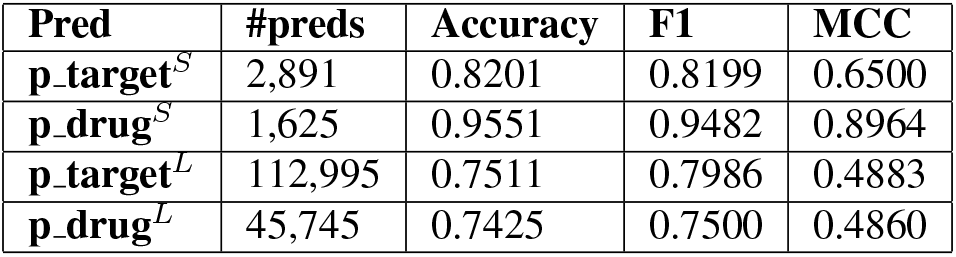
Results of sequence identity baseline, and number of interactions analyzed.

**Figure 5:**
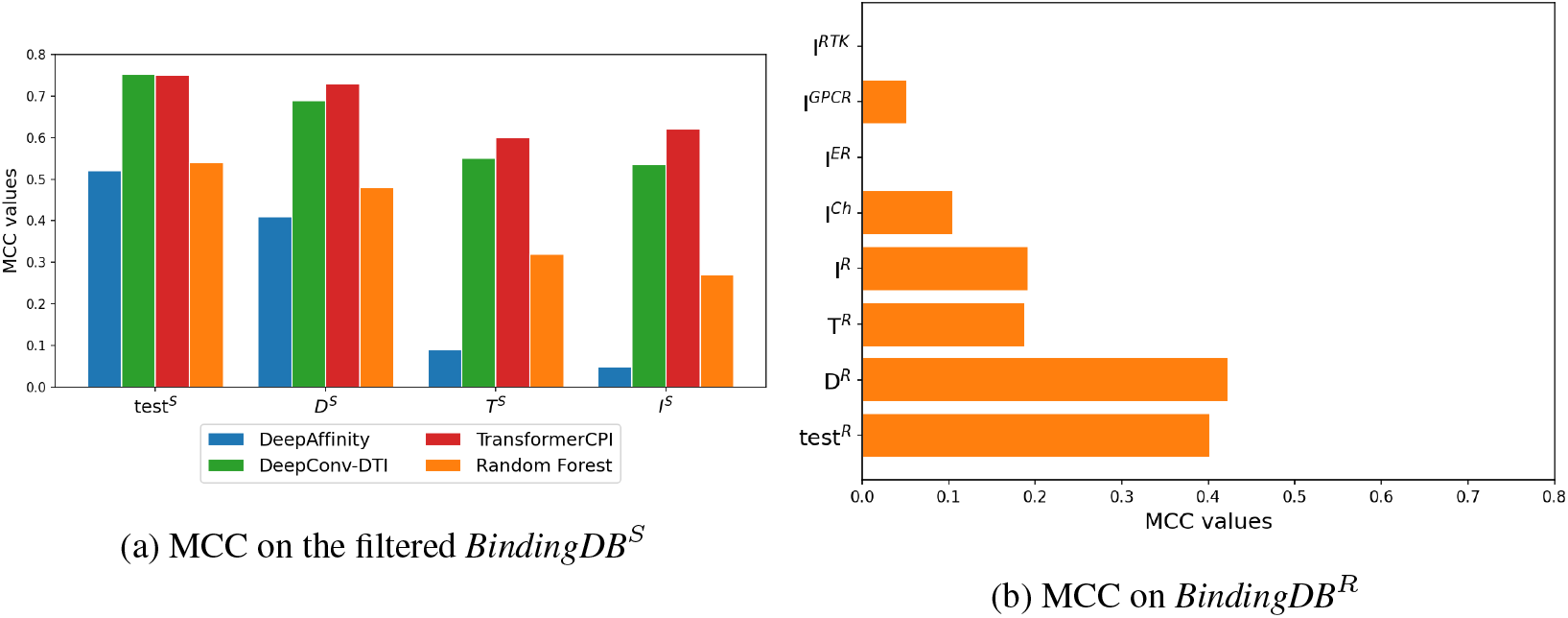
We derive 3 subsets for each benchmark dataset collecting any interaction where: (1) the drug is not in the training set (D), the target is not in the training set (*T*), neither the target nor the drug is in the training set (*I*). MCC observed on (a) the test set and subsets of *BindingDB^S^*, and on (b) the sets of *BindingDB^R^*. Figure (b) shows the MCC of a Random Forest.

**Figure 6:**
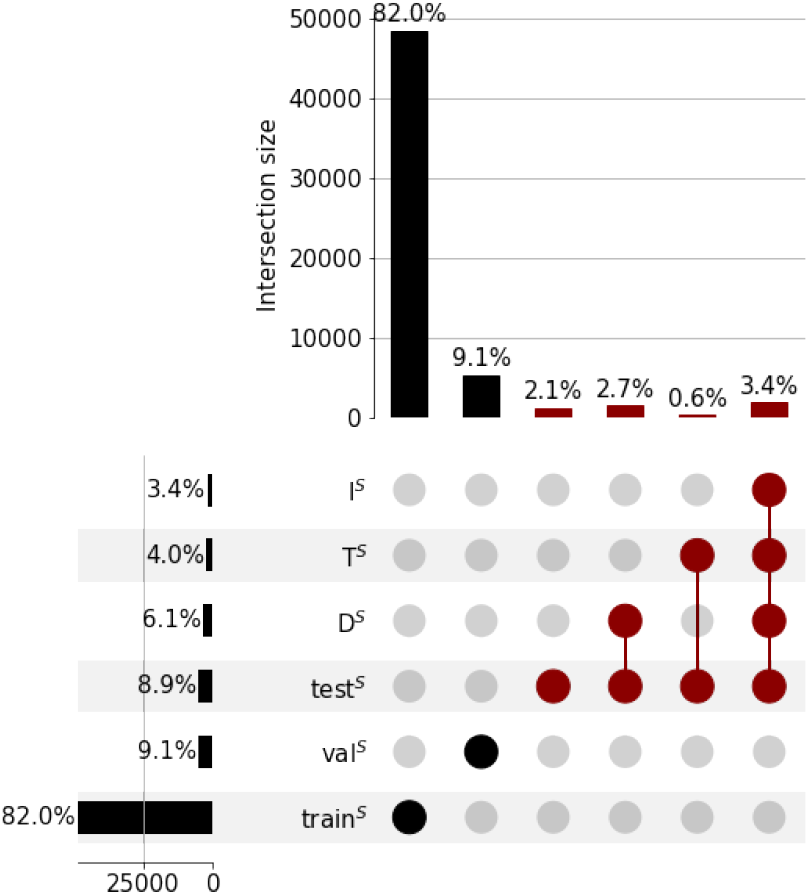
Overlaps of interactions in *BindingDB^S^* with respect to the entire benchmark dataset. The figure shows that there is no simple case of data leakage with the training set, in net contrast with what we observe on *BindingDB^L^* (see Figure 4a).

**Figure 7:**
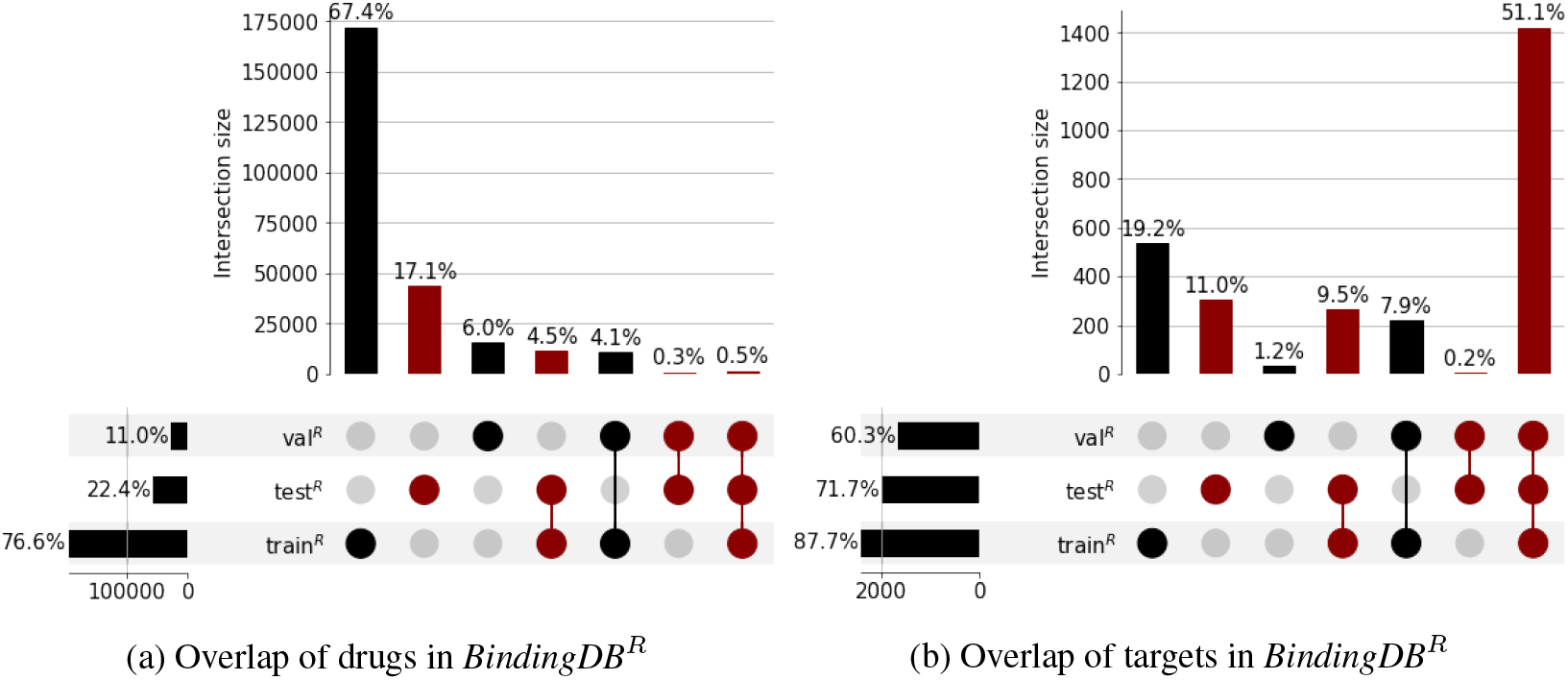
Overlaps of (a) drugs and (b) targets in *BindingDB^R^* with respect to the entire benchmark dataset. In both cases, the overlaps with the training set is reduced with respect to *BindingDB^L^* (see Table 1 and Figure 2b).

#### 3.3.2 BindingDB^L^

The large overlap between *train^L^* and *test^L^* leaves only 64 interactions in *I^L^*. Therefore, we create a new benchnmark dataset, called *BindingDB^R^*, by resplitting the training, validation, and test set of *BindingDB^L^* (see Table 2). To do so, we randomly select 10% of targets and 10% of drugs, and allocate any interaction involving those to *test^R^*. We allocate 90% of any other interaction to *train^R^*, and the remaining 10% to *val^R^*. Thus, all samples in *test^R^* are in *D^R^, T^R^*, or *I^R^*.

*BindingDB^R^* contains a similar ratio of samples as in *BindingDB^L^*, while reducing the overlap of both drugs and targets between training and test set, and remediating the data leakage in *test^L^* (see Figure 4a). We also filter the four OOD sets removing any interaction involving a drug or target in *train^R^*, i.e. *I^Ch^, I^ER^, I^GPCR^*, and *I^RTK^*.

We train a Random Forest on *train^R^* and we expect analogous results for the deep learning methods in this study. The Random Forest performs similarly well on *test^R^* and *D^R^*, and worse on *T^R^* and *I^R^*, as shown in Figure 5b. This matches what we observe for *BindingDB^S^* in Section 3.3.1, and provides additional evidence of the non-independence of drugs and targets within interactions.

The predictive performance on *T^R^* and *I^R^*, and to same extend those on *T^S^* and *I^S^*, resemble the results on *test^Ch^, test^ER^, test^GPCR^*, and *test^RTK^*. Thus, we see our approach as an alternative solution to evaluate OOD predictive performance, without depriving the training set of entire protein families (as done for the four OOD sets).

## 4 Conclusion

In this work, we perform an independent benchmark of recent DTI predictors based on deep learning. We find information leakage in previously used DTI benchmark datasets due to the non-independent nature of drugs and targets within interactions. We examine if such information leakage is related to overoptimistic predictive performance and relatively poor generalization observed. To support this idea, we show the high predictive performance obtained with a baseline analysis based solely on drugs and targets identity. To remove this source of bias and improve the assessment of DTI predictors, we propose a novel approach for assessing DTI predictors with respect to the content of their training set. Our novel approach results in a more comprehensive assessment of DTI predictors, discerning the OOD scenarios seen at test time, and highlighting potential source of bias without altering the training set. For future work, we aim to consider similarity of drugs and targets, instead of identity, to extend our minimal solution for evaluating the generalization capability of DTI predictors.

## Acknowledgments

The authors are grateful to Brian Kidd and Wilbert Copeland from BMS, and the anonymous reviewers for useful comments and suggestions.

## A Appendix

## Notes

### Competing Interest Statement

The authors have declared no competing interest.

